# Globally-deployed sorghum aphid resistance gene *RMES1* is vulnerable to biotype shifts but being bolstered by *RMES2*

**DOI:** 10.1101/2023.11.07.566092

**Authors:** Carl VanGessel, Brian Rice, Terry J. Felderhoff, Jean Rigaud Charles, Gael Pressoir, Vamsi Nalam, Geoffrey P. Morris

## Abstract

Durable host plant resistance (HPR) to insect pests is critical for sustainable agriculture. Natural variation exists for aphid HPR in sorghum (*Sorghum bicolor*) but the genetic architecture and phenotype has not been clarified for most sources. To assess the threat of a sorghum aphid (*Melanaphis sorghi*) biotype shift, we characterized the phenotype of *Resistance to Melanaphis sorghi 1* (*RMES1*) and contributing HPR architecture in globally-admixed populations selected under severe aphid infestation in Haiti. We found *RMES1* reduces sorghum aphid fecundity but not bird cherry-oat aphid (*Rhopalosiphum padi*) fecundity, suggesting a discriminant HPR response typical of gene-for-gene interaction. A second resistant gene, *RMES2*, were more frequent than *RMES1* resistant alleles in landraces and historic breeding lines. *RMES2* contributes early and mid-season aphid resistance in a segregating F_2_ population, however *RMES1* was only significant with mid-season fitness. In a fixed population with high aphid resistance, *RMES1* and *RMES2* were selected for demonstrating a lack of significant antagonistic pleiotropy. Associations with resistance co-located with cyanogenic glucoside biosynthesis genes support additional HPR sources. Globally, therefore, a vulnerable HPR source (*RMES1*) is bolstered by a second common source of resistance in breeding programs (*RMES2*) which may be staving off a biotype shift.

**HIGHLIGHT:** The globally-deployed sorghum aphid resistance gene, *RMES1*, reduces aphid reproduction and therefore is vulnerable to a biotype shift. A second major gene, *RMES2*, and cyanogenesis may increase global durability of resistance.

## INTRODUCTION

Plant breeding indirectly affects insect populations by applying selection pressure via deployed host plant resistance (HPR) which deters infestation. It is important for breeders to consider what HPR is being deployed in order to reduce the likelihood of population shifts in agronomically important pests or their emergence. Insect populations have regularly overcome HPR with genetic or geographical shifts into open niches. For example, agronomically important biotypes of Russian wheat aphid (*Diuraphis noxia*) and greenbug (*Schizaphis graminum*) have shifted in the Great Plains in the 20th century (Harris-Shultz *et al*., 2020). The *Nr* and *Ag1* resistance alleles were overcome by resistance breaking aphid biotypes of the lettuce root aphid (*Pemphigus bursarius*) and Large Raspberry aphid (*Amphorophora agathonica*), respectively, in Europe (Keep, 1989; Arendt *et al*., 1999). Climate change is expected to exacerbate this problem as climate-change-induced hybridization and habitable range expansion drive genetic diversity (Arce-Valdés and Sánchez-Guillén, 2022).

Aphids are economically significant pests which remove photoassimilates and vector viruses. Plants have multiple layers of HPR against aphids including morphological barriers and chemical compositions which can prevent feeding and infestation by deterring aphid behavior (antixenosis) and/or reducing fecundity (antibiosis) (Nalam *et al*., 2019). In contrast, tolerance is another type of plant response that allows the plant to maintain fitness under moderate infestation (Painter, 1951). However, our understanding of the molecular mechanisms behind aphid HPR in plants is limited. In one example of monogenic constitutive antixenosis, sorghum aphid (*Melanaphis sorghi*; Theobald 1904) feeding preference was affected in choice-assays by a *bloomless* gene knockout which lacked cuticular wax while reproduction in no-choice assays was not (Cardona *et al*., 2022). The cloned *Mi-1* and *Vat* encode nucleotide binding leucine-rich repeat (NLR) receptors (Nombela *et al*., 2003; Dogimont *et al*., 2014). Translational evidence from plant-pathogen research suggests that a gene-for-gene arms race could occur in plant-aphid systems, since resistance breaking biotypes of *Mi-1* and *Vat* have been identified and is one hypothesis for the boom and bust cycles of greenbug biotype-specific HPR (Kaloshian, 2004; Dogimont *et al*., 2010).

Sorghum (*Sorghum bicolor* L. [Moench]) is among the world’s most important cereals and a staple crop for small-holder farmers in semi-arid regions. A severe sorghum aphid outbreak occurred in the Americas beginning in 2013, believed to be due to a range expansion rather than biotype shift, and dispersed to all production regions within five years of its introduction (Armstrong *et al*., 2015; Harris-Shultz *et al*., 2020; Nibouche *et al*., 2021). One major HPR source to sorghum aphid is the globally-deployed, *<u>R</u>esistance to <u>Me</u>lanaphis <u>s</u>orghi <u>1</u>* (*RMES1*), on chromosome 6 (Chr06) found in African landraces where *M. sorghi* was first described (Vuillet, 1914; Muleta *et al*., 2022). *RMES1* is rare in global sorghum diversity, however it was swept to fixation in a recently established Haitian breeding population founded on globally admixed genotypes from East and West Africa (Muleta *et al*., 2022). A second sorghum aphid HPR QTL (*RMES2*) co-localized with a WRKY transcription factor whose functional allele may act as a regulatory hub for induced defenses (Poosapati *et al*., 2022). The prevalence and utility of this second sorghum aphid HPR source for breeding programs has not been established. In addition to cuticular waxes, other phytochemical herbivore deterrents such as the cyanogenic glucoside dhurrin have been proposed to contribute HPR(Woodhead *et al*., 1980; Poulton, 1990).

The durability of HPR to aphid populations is expected to vary between tolerance and resistance as they apply different selection pressures, but study systems to test this are not tractable (Peterson *et al*., 2017). The source of greenbug biotype C resistance in sorghum was considered tolerance but was overcome by biotype E shortly after wide deployment (Hackerott *et al*., 1969). A biotype C and E resistant grain sorghum breeding line, RTx2783, was identified as having tolerance and antibiosis-resistance to sorghum aphid (Armstrong *et al*., 2015). RTx2783 contains the *RMES1* allele and was donor for aphid resistance in many cultivars grown on the Great Plains (Muleta *et al*., 2022). Hypotheses on whether *RMES1* provides tolerance or resistance to sorghum aphid, and whether it provides discriminant or broad-spectrum resistance among Aphididae spp., have not been tested but would provide insight on its durability and the likelihood of biotype shifts.

Sorghum aphid and *S. bicolo*r is an agronomically relevant system to study HPR and pest population dynamics (Zapata et al., 2018). There are competing hypotheses on whether *RMES1* provides resistance (antibiosis or antixenosis) or tolerance and whether that phenotype extends to other aphid species. The contribution of *RMES2* to global breeding is unknown and molecular breeding tools have not been developed (Poosapati *et al*., 2022). It is also unclear whether *RMES1* and *RMES2* act additively or are epistatic. Here, we define the *M. sorghi*-discriminant resistant phenotype of *RMES1* and demonstrate the potential for *RMES2* to bolster *RMES1* resistance based on global germplasm. Finally, we confirm the contribution of both HPR sources in a breeding program where *RMES2* is being selected alongside *RMES1*.

## MATERIAL AND METHODS

### Near-isogenic line development

*RMES1* near-isogenic lines (NILs) were developed with a donor parent IRAT204 (*M. sorghi* resistant, *RMES1* donor) and recurrent backcrossing to RTx430 (*M. sorghi* susceptible). Single plant selections were made at the BC_x_F_2_ using a KASP marker for *RMES1* (Sbv3.1_06_2892438R, (Muleta *et al*., 2022) and homozygous plants were backcrossed to RTx430. BC_2_F_3_ progeny (NIL+ containing resistant *RMES1* allele, NIL-containing susceptible *RMES1* allele) and parental genotypes were used for aphid bioassays and whole genome resequencing.

### Whole genome resequencing of NILs

Genomic DNA was collected from leaf tissue of BC_2_F_3_ NIL+, NIL-, IRAT204, and RTx430. Four plants of each genotype were grown, DNA extracted with Zymo Plant DNA Isolation Kits, and sequenced individually. Samples were sequenced at the Genomics Shared Resource Core at the University of Colorado Anschutz Medical Campus. Raw reads were trimmed using trimmomatic v0.39 and mapped to the RTx430v2 reference (https://phytozome-next.jgi.doe.gov/info/SbicolorRTx430_v2_1) with BWA v0.7.17-r1188 (Li and Durbin, 2009). Duplicate reads were identified using Picard v2.26 (http://broadinstitute.github.io/picard) (McKenna *et al*., 2010). Finally, variants were called using GATK v4.2.5.0.

### Aphid cultures

*M. sorghi* used in this study were received from Dr. Scott Armstrong at the USDA-ARS Stillwater, Oklahoma. Aphids were reared on Tx7000 seedlings under laboratory conditions as previously described (Nalam et al. 2021). Seedlings were grown in 4.5” pots with potting soil and top layer of sand to reduce damping off at a temperature of 24 ± 1°C and a photoperiod of 16:8 (L:D) hours. Colonies were housed in cages covered with insect-proof cloth. Bird cherry-oat aphid (*Rhopalosiphum padi*) cultures were collected from the CSU greenhouses and maintained on Tx7000 seedlings similar to *M. sorghi* cultures.

### Choice assay for aphid settling preference

A choice assay was done to determine aphid settling preference at the seedling stage. A pairwise comparison was done with NIL+ and NIL-. Seedlings of each genotype were grown approx. 2 inches apart in 1 gallon pots using potting soil and a top layer of greens grade. Seedlings were thinned to one plant of each genotype per pot. At 3-4 weeks of age, or 2-3 leaf stage, twenty 3-4 day old apterous *M. sorghi* were placed in the center of a paper bridge between the seedlings. A clear plastic cylinder was placed over the plants to prevent aphids from leaving the pot with an organdy cloth covering for ventilation. The number of aphids on each plant were counted at 6, 12, 24, and 48 hours post infestation (hpi). Statistical analyses were done in R v4.2.2 to detect differences in *M. sorghi* preferences between genotypes. A Student’s *t*-test was used to compare genotypes at each time point.

### No-choice assay for aphid fecundity

No-choice assays were used to compare aphid fecundity on various genotypes. A single seedling was grown in 6 inch pots using potting soil and a top layer of greens grade. At 3-4 weeks of age, three 3-4 day old apterous aphids were placed at the base of the seedlings with a camel hair brush. A clear plastic cylinder was placed over the plant to prevent aphids from leaving the pot with an organdy cloth covering for ventilation. The number of aphids on each plant were counted daily for a week at ∼12 PM. A no-choice assay with *R. padi* was used to determine cross-resistance. NIL+ and NIL-lines were infested with three 4-5 day old apterous aphids and counted daily for one week and at two weeks after infestation.

### Resequencing-based GWAS of community association panels

Whole genome resequencing for 665 sorghum genotypes from the Sorghum Association Panel (SAP) and Bioenergy Association Panel (BAP) were used for association analyses. Raw reads for the SAP were retrieved from the European Nucleotide Archive (RJEB50066) (Boatwright *et al*., 2022). Raw reads for the BAP were retrieved from DRYAD (https://doi.org/10.5061/dryad.4b8gtht99) (LeBauer *et al*., 2020).

Raw reads for both association panels were mapped to BTx623 v5.1 (sorghumbase.org/Sorghum_bicolorv5) reference using BWA-mem (Li and Durbin, 2009). Samtools (Li *et al*., 2009) was used to select properly paired reads and sort, Picard v2.26 (http://broadinstitute.github.io/picard) was used to remove duplicate reads, Samtools was used to remove low quality reads (“-Q 30”), and VarScan (https://sourceforge.net/projects/varscan/) used for variant calling. Variants were filtered using bcftools v1.15.1 commands ‘F_MISSING < 0.9’ and ‘MAF > 0.01’ (Danecek *et al*., 2021). Variants were imputed with BEAGLE v5.2 using default parameters (Browning *et al*., 2018). The principal components (PC) of a random subset of 500,000 SNP variants was used to estimate population structure computed by TASSEL 5.0 CLI -PrincipalComponentsPlugin function. A general linear model was used to determine associations with the -FixedEffectLMPlugin using the first three PCs and normalized sorghum aphid phenotypes (Bradbury *et al*., 2007; Poosapati *et al*., 2022). A second association analysis was performed with the highest associated variant at *RMES2* (S09_61521444) included as a fixed-effect covariate. Manhattan plots were generated in base R (v4.2.2).

The most recent reference genome version (BTx623v5) is used throughout this paper and coordinates refer to the v5 coordinate system unless otherwise noted. For example S09_61433682 refers to the variant at 61,433,682 bp on Chr09 in the BTx623v5 genome while S09_57630053.v3 represents a variant at 57,630,053 in BTx623v3 and S06_3222901vRTx430 represents a variant at 3,222,901 on the RTx430v2.1 with a different coordinate system.

### Population genomic analyses of *RMES2* in landraces and early breeding germplasm

Unimputed allele calls from the resequenced lines for Chr06_3096975 and Chr09_61521444 for 268 sorghum accessions with longitude, latitude and/or germplasm origin was used to determine allelic distributions for *RMES1* and *RMES2* (Supplemental Data 5). Metadata was collected from GeneSys (www.genesys-pgr.org/c/sorghum). Geographic distributions and pie charts were generated using r/ggplot2 (v3.4.1).

### Fixation analysis of globally-admixed germplasm resistant to sorghum aphid

The fixation analysis used here was previously described (Muleta *et al*., 2022). Briefly, the Chibas sorghum breeding program was founded in 2013 using global germplasm and extensively intercrossed using the *ms3* sterility system. Each generation, 5%-10% of the population is selected for grain yield potential, maturity, and height preferences of Haitian farmers. Selected females are pollinated by five randomly selected male lines (of those selected for generation advancement) and the next generation begins again. Low-input conditions which approximate small-holder farmer growing practices were used for population development allowing for heavy sorghum aphid infestation in year-round growing seasons with tropical environments. The resulting aphid-resistant admixed population (N=296) was genotyped using genotype-by-sequencing and compared to a separate set of accessions representing global diversity (N=767) (Morris *et al*., 2013; Lasky *et al*., 2015). Pairwise SNP differentiation (*F*_ST_) was determined between admixed breeding lines and global accessions and outlier loci detected based on inferred distribution of neutral *F*_ST_.

### Association analysis of globally-admixed F_2_ population segregating for aphid resistance

A second F_2_ population was developed by the Chibas sorghum breeding program and used for genetic trait dissection. Lines from the globally-admixed germplasm resistant to sorghum aphid (fixed for *RMES1*) were crossed to new diverse material in order to develop globally-admixed F_2_ lines segregating for aphid resistance phenotypes (Muleta *et al*., 2022). Phenotyping data was collected in spring-summer 2022 in Haiti under heavy sorghum aphid infestation. Susceptible checks had over 1,000 aphids per plant, infesting all leaves, and covering ⅓ to ½ of leaf area. Fields were irrigated once a week and no pesticides or fertilizers were used. Plants were phenotyped for fitness (alive/dead) at flowering initiation (6-7 leaf) and booting (8-9 leaf) growth stages.

Tissue was collected for genotyping approximately one month after planting and genotyped using Diverse Array Technology (Jaccoud *et al*., 2001). Sequencing was mapped to the BTx623v3.1 reference genome (McCormick *et al*., 2018). Marker positions were converted to the BTx623v5.1 coordinate system by BLAST (https://phytozome-next.jgi.doe.gov/) with 100 bp flanking segments. Data was processed in R package dartR, and markers which were monomorphic, had <50% call rate, or <90% repeatability were removed. Data was converted to VCF format using a custom R script and imputation was done using Beagle 5.4 (Browning *et al*., 2018). Data was converted to numeric representation of reference alleles (0,1,2). After filtering, there were 1,172 individuals with 8,195 markers.

Fitness was modeled as a binary phenotype using the generalized linear model (1);

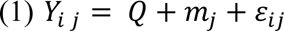

where *Y*_*i*_ is the observed phenotype state for plant *i* (coded as 0 for dead and 1 for alive) and fit with a binomial logit function; *G*_*i*_ is the random effect of the i^th^ plant distributed 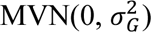 where 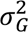 is the genetic variance, and *ε_i_* is the residual error; *Q* is a matrix of *n by two* fixed effects accounting for population structure estimated from the first two principal components of the genome-wide SNP set; *m* is the fixed effect of marker *j*. A custom script was used to run model (1) with AsremlR for each marker (Butler *et al*., 2017).

## RESULTS

### Marker-assisted selection isolates *RMES1* in near-isogenic lines

Testing hypotheses on specific HPR sources such as *RMES1* and its phenotype requires suitable germplasm which isolates resistant and susceptible alleles such as near-isogenic lines (Figure 1a). We therefore backcrossed *RMES1*-donor IRAT204 to the susceptible RTx430 and generated BC_2_F_3-4_ lines. Whole-genome sequencing confirmed the marker-assisted backcrossing successfully reduced the donor genome complement in NILs. There was 71.7% and 24.8% of the NIL+ genome fixed for the recurrent and donor parent genomes, respectively, and the remaining 3.4% was segregating (Figure 1b). The NIL-genome was 80.7% and 14.2% fixed for the recurrent and donor parent genomes, respectively, and 5.1% segregated (Figure 1b). Comparing between NIL+ and NIL-sibling lines, 83.6% of the genome was isogenic and the remaining 16.3% was segregating, including the majority of Chr06 (Figure 1c). In the region of *RMES1* on Chr06, both NIL lines were fixed - however, there is a breakpoint in NIL+ lines between 2,051,868 bp and 2,169,917 bp (RTx430v2 coordinates). The *RMES1* region, mapped in BTx623, corresponds to 2.9–3.1 Mb of the RTx430v2 genome and is therefore within the introgression. The previously reported *RMES2* SNP (S9_57630053v3) is monomorphic in IRAT204 and RTx430 as well as falling ∼5 Mb outside of the introgressed region on Chr09. Therefore, these NILs allow us to test phenotypic hypotheses regarding *RMES1* without confounding sources of additional HPR.

**Figure 1.**
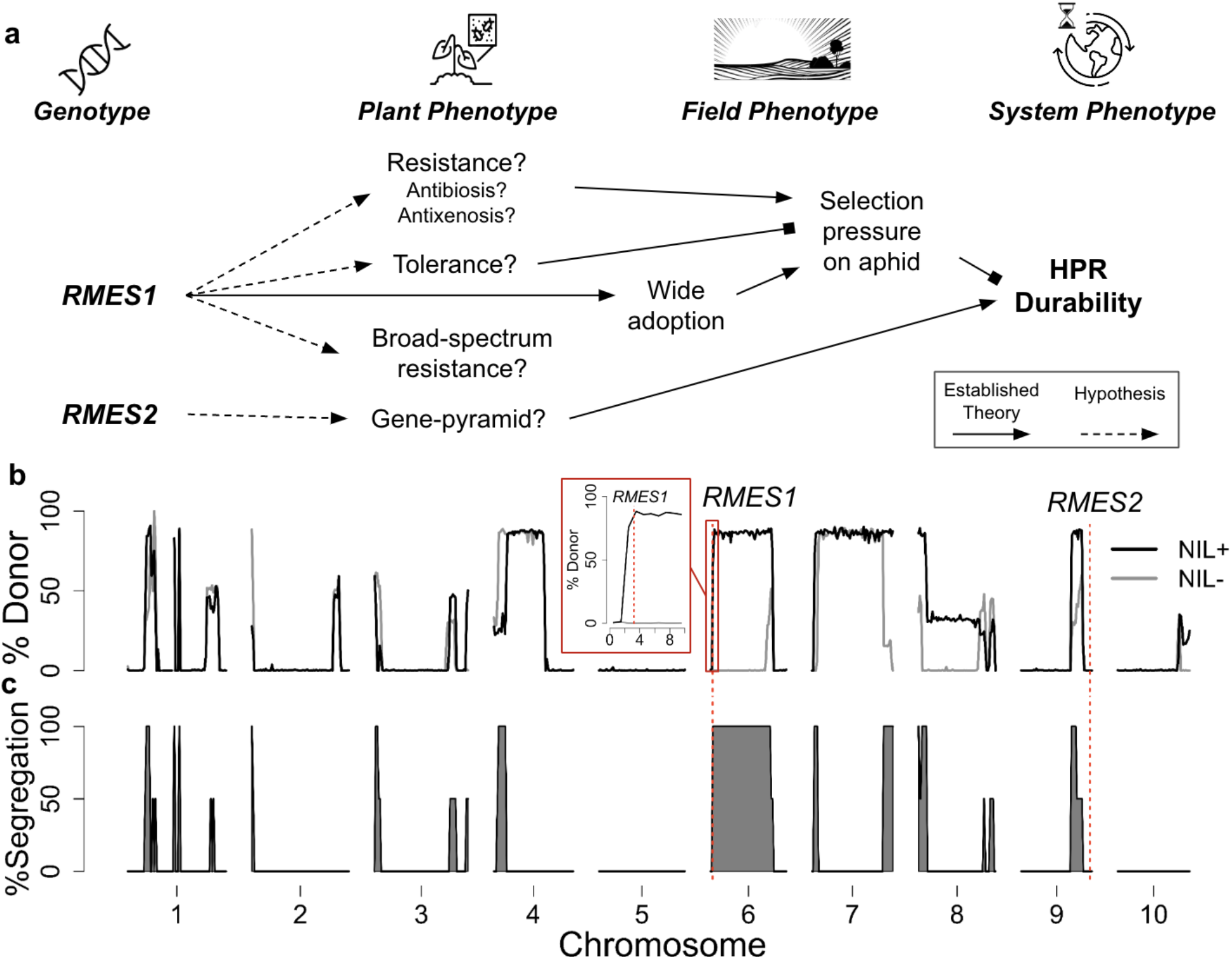
Genotype to phenotype hypotheses on *RMES1* to be tested with near-isogenic lines. a) The genotype to phenotype map for *RMES1, RMES2,* and host plant resistance (HPR) durability containing several established and hypothesized relationships. b) Donor (IRAT204, 100%) genome contribution in BC_2_F_3_ NILs relative to recurrent genome (RTx430, 0%) using whole-genome sequencing mapped to RTx430v2.1 reference genome. The inset panel is a close-up of the *RMES1* region on chromosome 6. c) Genomic regions segregating between NILs determined from WGS. Red dotted line indicates *RMES1 F*_ST_-identified SNP on chromosome 6 (S06_2995581v3 = S06_3222901vRTx430) (Muleta *et al*., 2022) and the original *RMES2* GWAS-identified SNP on chromosome 9 (BTx623v3.1 S9_57630053, RTx430v2.1 S09_54801196) (Poosapati *et al*., 2022). Genomic analyses indicates that, as intended, NILs segregate for *RMES1* but not *RMES2*.

### *RMES1* reduces fecundity of sorghum aphid

To determine whether antibiosis is a component of *RMES1*, we assessed aphid reproduction in a no-choice assay. Differences in population growth would indicate the *RMES1* mechanism is retarding infestation and placing selection pressure on aphids. The number of aphids was lower on NIL+ than NIL-at 7 days post infestation (7 dpi) (*P* < 0.001; Figure 2b). A moderately significant difference (*P* = 0.025) was first seen at 3 dpi and highly significant (*P* = 0.001) at 6 dpi. The initial infestation of three aphids grew to an average of 28 ± 4 aphids and 53 ± 4 aphids on NIL+ and NIL-, respectively.

**Figure 2.**
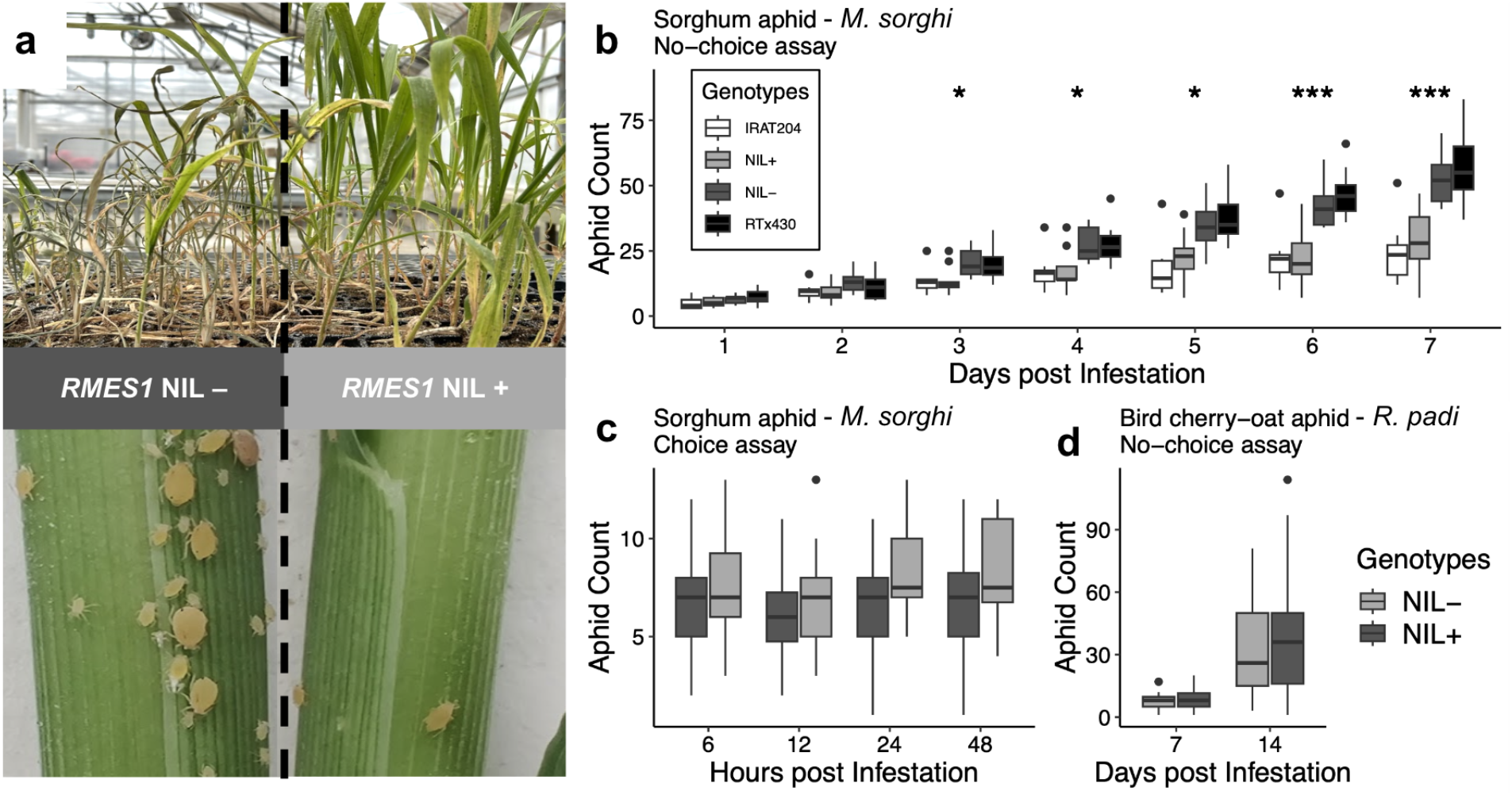
*RMES1* provides antibiosis-resistance to sorghum aphid but not all cereal aphids. a) Representative photographs of *M. sorghii* aphid infestation on *RMES1* NILs. b) No-choice assay of sorghum aphid counts over 7 day infestation on NILs and parent genotypes (*N* = 9). c) Choice assay of sorghum aphid counts over 48 hour infestation on NILs (*N* = 16). d) No-choice assay of *R. padi* at 7 dpi and 14 dpi (*N* = 18). Significant pairwise comparison between NILs (Student’s *t*-test) shown with asterisks (*P* < 0.05 *, *P* < 0.01 **, *P* < 0.001 ***)

Antibiosis affects the behavior and settling preference of aphids and was tested for using a choice assay. We found that aphid settling was not significantly different between NILs at any time point in the first 48 hours of infestation (Figure 2c). This indicates that sorghum aphid feeding choice is not determined by *RMES1* and suggests a lack of constitutively expressed epidermal or volatile features that deter aphid settling. While tolerance may be an additional component of *RMES1* phenotype, the significant antibiosis component is placing selection pressure on aphid populations.

### *RMES1* does not provide resistance to bird cherry-oat aphid

The presence of *RMES1* in the commonly used breeding line RTx2783 with greenbug resistance led to the hypothesis that the locus provides broad-spectrum resistance across Aphididae species (Armstrong *et al*., 2015). A no-choice assay of *R. padi* on *RMES1* NILs showed no effect on reproduction over 7 and 14 day infestation (*P* = 0.3; Figure 2d). Aphid reproduction was lower for *R. padi* than *M. sorghi* with aphids populations growing from 3 to an average 8 ± 1 aphids after one week infestation and average 30 ± 9 aphids after two weeks. This indicates *RMES1* is at least partially discriminant in providing Aphididae resistance, although it may still play a role in greenbug resistance.

### GWAS with resequencing confirms that *RMES1* resistance allele is rare in global landraces

It was previously shown that the *RMES1*-associated SNP with the highest fixation signature in a Haitian breeding program (S06_2995581v3) was rare in a global diversity panel. Having used GBS data and a breeding population to identify this *RMES1* associated SNP, it remains possible that a SNP in higher LD with *RMES1* exists and is at detectable frequency in association panels. We retested the rare *RMES1* hypothesis by combining recently published phenotypes for sorghum aphid resistance in global panels of landraces and improved lines (SAP and BAP) with whole-genome resequencing data and the *S. bicolor* BTx623v5.1 reference and performed genome-wide association analyses. We did not observe a peak at *RMES1* on Chr06 (Figure 3a, Supplemental Data 1). We included the major association for resistance, S09_61521444, as a fixed-effect covariate in our GLM model and confirmed the Chr06 locus was not significant after controlling for potentially confounding variation (Figure 3b, Supplemental Data 2). An association at S01_1536310 was 358 kb from the cyanogenic biosynthetic gene cluster, between 1.07 Mb and 1.18 Mb on Chr01 (Figure 3b) (Hayes *et al*., 2015). Additional loci on chromosomes 2, 3, and 10 were apparent when the major Chr09 HPR was accounted for and are candidates for additional quantitative HPR sources (Table 1).

**Figure 3.**
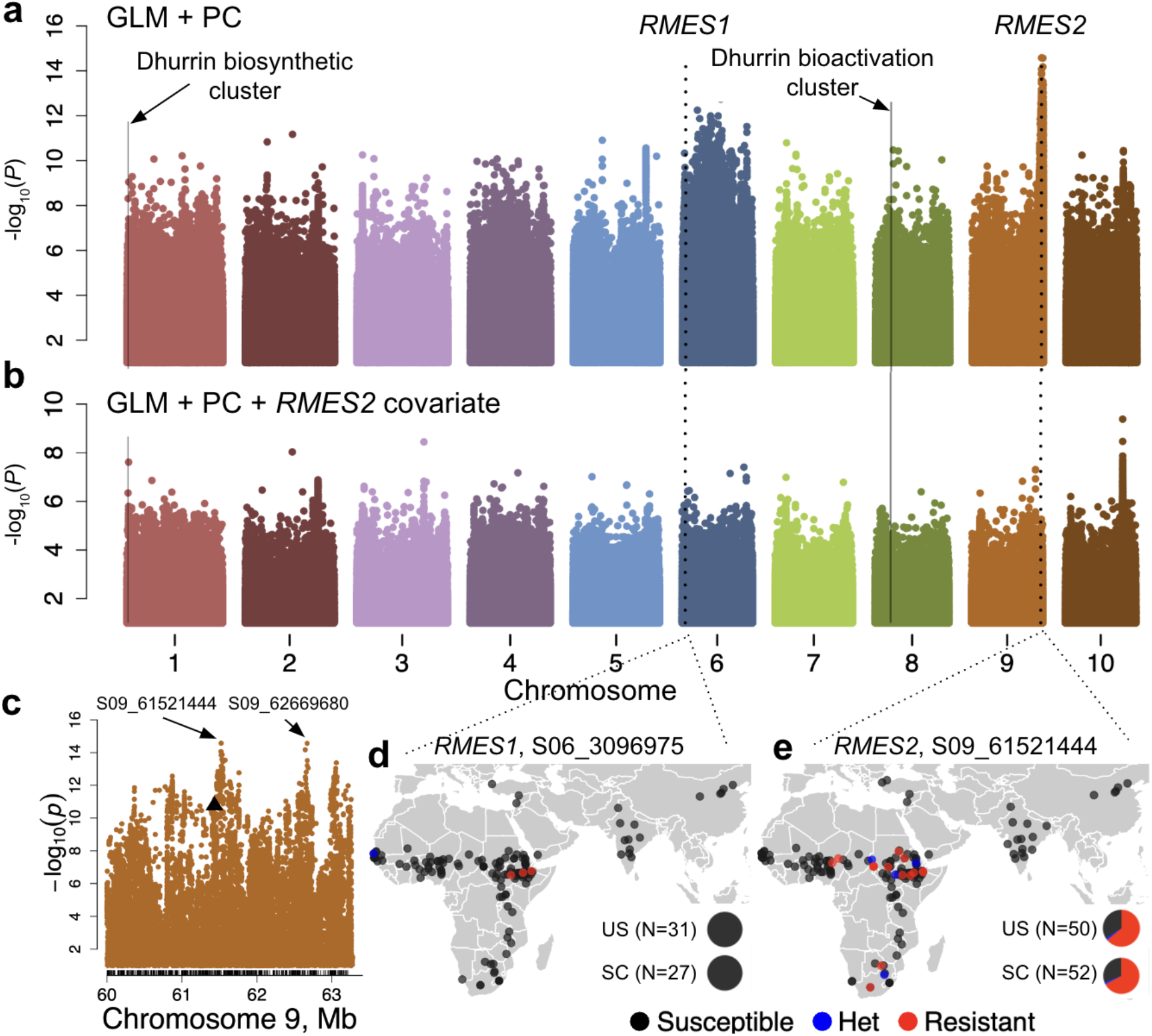
Genome-wide associations with *M. sorghi* resistance in global diversity panels show *RMES1* resistance is globally rare while *RMES2* is globally common. a) Association of resequencing variants determined with a general linear model (GLM) that included principal components (PC) 1-3 as fixed effects, phenotypes from Poosapati *et al*., 2022. b) Association of resequencing variants determined with a GLM that included principal components 1-3 of population structure and the peak *RMES2* SNP from the GLM (S09_61521444) as fixed effect covariate. c) Chr09 associations at *RMES2* highlighting the two most significant SNPs from the GLM + PC. The previously reported variant in the *SbWRKY86* gene model (S09_57630053v3) is indicated by the black triangle. Black lines below the associations indicate gene models in the region. d) Distribution of *RMES1*-associated SNP previously identified (Muleta *et al*., 2022). e) Distribution of *RMES2*-associated SNP identified in the present study. Resequencing data available for historic breeding germplasm (US) and lines from the sorghum conversion program (SC) was used to estimate the frequency of alleles in foundational US germplasm.

**Table 1:** Loci with significant associations with resistance (corresponding to Figure 3, Figure 4b,c)

| Marker | Chr | Pos (BTx623v5.1) | *P* value | GWAS Analysis |
| --- | --- | --- | --- | --- |
| S09_61521444 | 9 | 61521444 | 2.64E-15 | SAP+BAP |
| S09_62669680 | 9 | 62669680 | 2.71E-15 | SAP+BAP |
| S10_50970852 | 10 | 50970852 | 4.13E-10 | SAP+BAP, *RMES2* fixed covariate |
| S03_59780623 | 3 | 59780623 | 3.58E-09 | SAP+BAP, *RMES2* fixed covariate |
| S02_41893124 | 2 | 41893124 | 9.24E-09 | SAP+BAP, *RMES2* fixed covariate |
| S01_1536310 | 1 | 1536310 | 2.43E-08 | SAP+BAP, *RMES2* fixed covariate |
| S09_61988551 | 9 | 61988551 | 8.76E-13 | Segregating F2, mid-season survival |
| S06_2170466 | 6 | 2170466 | 8.33E-12 | Segregating F2, mid-season survival |
| S08_48830246 | 8 | 48830246 | 6.00E-13 | Segregating F2, mid-season survival |
| S08_11769483 | 8 | 11769483 | 2.46E-14 | Segregating F2, early-season survival |
| S01_78081631 | 1 | 78081631 | 5.00E-16 | Segregating F2, early-season survival |
| S03_66114610 | 3 | 66114610 | 1.53E-13 | Segregating F2, early-season survival |
| S09_59794306 | 9 | 59794306 | 5.54E-13 | Segregating F2, early-season survival |
| S01_1253672 | 1 | 1253672 | 1.01E-12 | Segregating F2, early-season survival |

### *RMES2* resistance allele is common in global diversity lines

*RMES2* was previously reported as a source of HPR in the combined association panel and *SbWRKY86* was proposed as the causal gene due to the strongest associations falling within the gene model (Poosapati *et al*., 2022). In order to test hypotheses on *RMES2* genomic position and inform marker development, we used resequencing data which contains ∼200 times more variants than previous GBS datasets and generated new genome-wide associations to reanalyze Chr09. We found the peak association (*P* < 10^-14^) at S09_61521444 (Figure 3a) for the resistant associated reference allele (T) and present in 213 of 665 genotypes (32%). The second strongest association at *RMES2* was found 1.1 Mb from the peak association at S09_62669680 (*P* < 10^-14^) (Figure 3c, Supplemental Data 1). The previously reported peak association (S09_57630053v3 / S09_61433682v5) located in the 3’ UTR of *SbWRKY86* (Sobic.009G238200) was 89 kb from the peak association and remained significant (*P* < 10^-10^) (Figure 3c) (Poosapati *et al*., 2022).

The *RMES1* allele identified via selection signatures in resistant breeding lines was previously shown to be globally rare and unique to East Africa (Muleta *et al*., 2022). The presence of the resistant *RMES2* allele (S09_61521444) in 32% of association panel genotypes suggested it was globally common relative to *RMES1*. We used resequencing data for genotypes with known landrace or breeding origin to test this hypothesis. Of 145 sorghum landraces, the resistant *RMES1* (S06_3096975) was present in just three (2.1%) while *RMES2* allele was present in 13 (9.0%) lines (Figure 3d,e, Supplemental Data 3). The geographic distribution of *RMES2* was broader than *RMES1* across Africa, however the highest concentration was in East Africa. The resistant *RMES2* allele was present at higher frequency in historic US breeding lines (67.3%) and sorghum conversion lines (34.0%) than landraces, whereas the resistant *RMES1* allele was absent in both historic sets.

### Both *RMES1* and *RMES2* are being selected in a globally-admixed breeding population

The prevalence of *RMES2* in global germplasm and association with aphid resistance supports its availability and value for breeding programs. The fixation scan previously reported on the phenotypically resistant admixed breeding germplasm was reanalysed in light of *RMES2* (Muleta *et al*., 2022). We found a strongly selected variant (S09_61842248, *P* < 10^-10^) at *RMES2* and ∼321 kb from the peak association panel variant S09_61521444 (Figure 4a). While *RMES2* variants were significant, the most significant variant on Chr09 was at S9_53496194 (*p* < 10^-14^) and 8 Mb from *RMES2*. In addition to *RMES1* and *RMES2*, outliers of fixation signatures colocated with staygreen (*Stg3a*, S2_60183555) QTL and 1.6Mb from the dhurrin bioactivation gene cluster (S08_11803397) (Table 2). These loci could represent additional traits under selection in the breeding program (e.g. post-flowering drought tolerance and forage quality), however they may also play a role in resilience to sorghum aphid.

**Figure 4.**
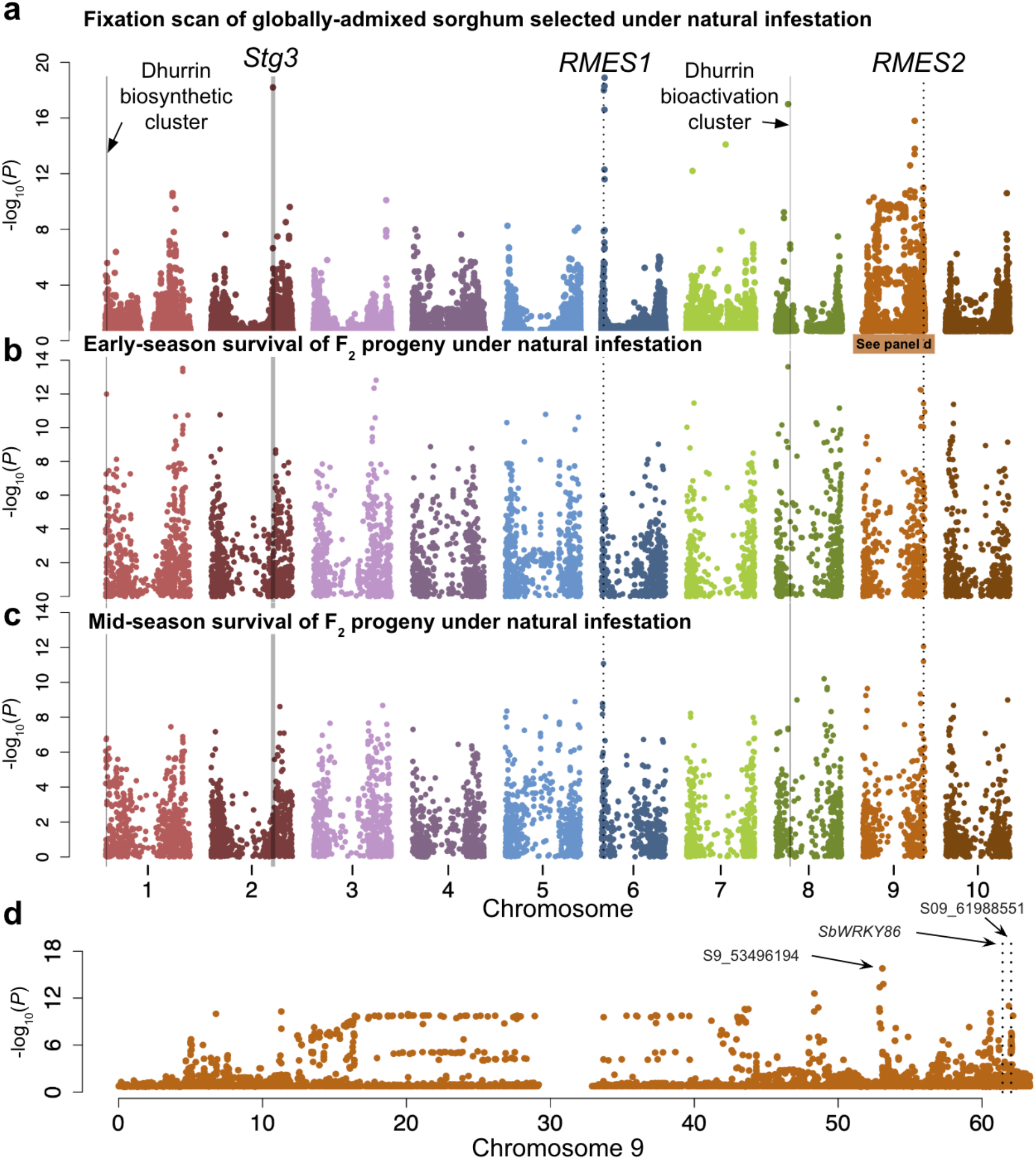
Both *RMES1* and *RMES2* contribute to aphid resistance in a globally-admixed breeding population. a) Fixation scores (*F*_ST_) determined for the phenotypically resistant admixed germplasm compared with the global diversity panel. Data replotted from (Muleta *et al*., 2022). Notable signatures are indicated with shaded lines (dhurrin biosynthesis and activation, *Stg3a* QTL region) (Cicek and Esen, 1998; Sanchez *et al*., 2002) or dashed lines (*RMES1* and *RMES2*). b-c) Associations with resistance under heavy aphid infestation in an F_2_ population of admixed germplasm segregating for resistance. b) Survival at flowering initiation associations. c) Survival at booting associations. The two strongest associations at booting for *RMES1* and *RMES2* are indicated by black dotted lines for all panels. d) *F*_ST_ scores on Chr09. S06_3096975 and S09_61988551 are used to indicate *RMES1* and *RMES2* in panels a-c.

**Table 2:** Loci with significant fixation signatures in globally-admixed breeding population (corresponding to Figure 4a)

| Marker | Chr | Pos (BTx623v5.1) |
| --- | --- | --- |
| S6_3096975 | 6 | 3096975 |
| S6_2993832 | 6 | 2993832 |
| S2_60183555 | 2 | 60183555 |
| S8_11803397 | 8 | 11803397 |
| S9_53496194 | 9 | 53496194 |
| S7_51523431 | 7 | 51523431 |
| S9_61842248 | 9 | 61842248 |

### *RMES1* and *RMES2* provide resistance in segregating F_2_ population founded on globally-admixed genotypes

*RMES1* was mapped in biparental populations for aphid resistance and through fixations signatures in the aphid resistant admixed germplasm, whereas *RMES2* has been mapped in association panels for aphid resistance (Wang *et al*., 2013; Muleta *et al*., 2022; Poosapati *et al*., 2022). We used a highly recombinant F_2_ population derived from crosses between resistant admixed genotypes and African landraces that was phenotyped for survival under heavy natural infestation of *M. sorghi* in order to test hypotheses on global genetic architecture contributing aphid resistance at early-season (flowering initiation) and mid-season (booting).

Despite expecting both *RMES1* and *RMES2* to contribute resistance throughout all growth stages, only *RMES2* (S09_59794306, *P* < 10^-12^) was associated with resistance early in development (Figure 4b, Supplemental Data 4). An association on Chr01 (S01_1253672, *P* < 10^-11^) was 76 kb from the dhurrin biosynthetic cluster while an association on Chr08 (S08_11769483, *P* < 10^-13^) was 1.6 Mb from the dhurrin bioactivation cluster. Other early-season resistance associations on Chr01 (S01_78081631, *P* < 10^-13^) and Chr03 (S03_66114610, *P* < 10^-12^) indicate HPR is polygenic at early growth stages. In contrast to early-season, the two strongest mid-season associations with resistance were found at *RMES2* (S09_61988551, *P* < 10^-11^) and *RMES1* (S06_2170466, *P* < 10^-11^) (Figure 4c, Supplemental Data 5). There was not a significant interaction between the two loci indicating no epistasis detectable in this population. The third most significant association at booting was on Chr08 at S08_48830246 (*P* < 10^-10^). Notably, *RMES2* was the only QTL which associated with resistance at both growth stages and there was no co-localization of dhurrin related QTL with resistance associations at the later growth stage.

## DISCUSSION

Here, we characterized the major source of HPR in a sorghum breeders toolkit (*RMES1*) and showed evidence that additional sources (*RMES2*, dhurrin biosynthesis QTL) are complementing the genetic architecture underlying global sorghum aphid resistance. We found that *RMES1* HPR provides antibiosis-resistance to sorghum aphid but not all Aphididae species. We next determined a strong likelihood that *RMES2* is present in breeding programs. Finally, we describe the collective contribution of *RMES1* and *RMES2* in a Haitian breeding program where breeding for sorghum aphid resistance is a priority. Co-localizations of aphid resistance associations with dhurrin pathway genes implicate cyanogenic potential as a quantitative source of resistance.

### Isolating *RMES1* in NILs allowed the phenotype and threat of biotype shift to be defined

Breeding focused research on HPR often identifies genetically dissimilar germplasm differing for traits such as antibiosis or tolerance which prevents mechanisms from being resolved at the genetic architecture level. It is important to mendelize traits using NILs, induced mutation, or gene-editing in order to test gene-specific hypotheses which untangle the many levels of HPR and elucidate individual mechanisms. The majority isogenic background and single resistance locus segregating in our NILs allowed us to define *RMES1* antibiosis-resistance for sorghum aphid (Muleta *et al*., 2022). Future investigations of the molecular mechanism will benefit from continued population development of these NILs.

The common breeding line RTx2783 contains the resistant allele of *RMES1* and showed tolerance and resistance to *M. sorghi* and *S. graminum* leading to the hypothesis that *RMES1* is pleiotropic for these traits (Armstrong *et al*., 2015). NILs had no difference in the *R. padi* no-choice assay (Figure 2d) showing that the molecular mechanism of resistance is not shared by all sorghum – Aphididae systems. Resistance to *S. graminum* has not been associated with the *RMES1* region in mapping studies supporting RTx2783 greenbug resistance being derived from non-*RMES1* sources (Harris-Shultz *et al*., 2020). *RMES1* is the only source of HPR with breeder-friendly marker technology and known to be used in public and private sorghum breeding programs (Muleta *et al*., 2022).

The molecular mechanism of *RMES1* remains to be tested but two hypotheses have been put forward. One candidate is *SbCAS1* (Sobic.006G016900), a *β*-cyanoalanine synthase gene involved in detoxification of HCN produced by the cyanogenic glucoside dhurrin (Gleadow *et al*., 2021). Cyanogenesis is involved in antibiosis-resistance to the Fall armyworm (*Spodoptera frugiperda*) in sorghum, however involvement in aphid resistance has not been tested (Gruss *et al*., 2022). A change in HCN detoxification strategy could have autotoxic consequences for the host, and the capacity to detoxify HCN is unlikely to vary significantly among cereal aphids i.e. *R. padi* and *M. sorghi* (Poulton, 1990). The other set of candidates involve a cluster of nucleotide-binding leucine-rich repeat (NLR) receptors, which recognize molecular patterns of infestation and activate host defenses, and are predicted in the region of *RMES1* in the BTx623 (susceptible) and RTx2783 (resistant) genomes (Wang *et al*., 2021; Muleta *et al*., 2022). These are strong candidates for causal genes due to numerous reports of NLR genes driving resistance in aphids and other pest systems (Dogimont *et al*., 2014; Snoeck *et al*., 2022; Boissot, 2023). This class of resistance mechanisms is expected to be less durable if the selection pressure is high and the herbivore-associated molecular pattern (HAMP) can withstand mutations to evade the host receptor. Such ‘gene-for-gene’ dynamics agree with the Aphididae-discriminant phenotype with *M. sorghi* and *R.padi*, and could lead to boom-and-bust cycles similar to those seen in cereal-biotic pest systems (Flor, 1971; Dogimont *et al*., 2010; Mundt, 2018). One or both cyanogenesis and NLR mechanisms could underlie *RMES1*.

As future studies establish the relative durability of antibiosis, antixenosis, and tolerance mechanisms, knowledge of *RMES1*-antibiosis will inform how to combine and utilize all HPR available. RTx2783 appears to harbor additional tolerance and HPR sources as it was reported to retain growth despite moderate aphid infestation (Limaje *et al*., 2018). Alternatively, pleiotropic interactions from *RMES1* may contribute additional tolerance mechanisms in different backgrounds. Regardless of the pleiotropic tolerance hypothesis, a significant reduction in fecundity of aphids on *RMES1* NILs demonstrates a selection pressure on *M. sorghi* and increasing the likelihood of a biotype shift. This HPR source should not be solely relied upon for crop protection as there is precedent for monogenic HPR breakdown in cereal-aphid systems.

### *RMES2* is moderately common in global landraces and contributes HPR in a diversity panel

High-quality resistance phenotyping of two association panels in conjunction with dense genotyping provided the opportunity to test hypotheses on architecture contributing resistance while also informing *RMES2* marker development which will provide a second breeder-friendly tool. The two strongest associations with GBS SNPs were inside the gene model of *SbWRKY86*, a transcription factor responsive to aphid infestation (Kiani and Szczepaniec, 2018; Poosapati *et al*., 2022). We found two regions 87.8 kb and 1.24 Mb from *SbWRKY86* which had a higher association than the original SNP in our analysis (Figure 3a). One methodological hypothesis for discrepancies is differences between the previous panel of 697 genotypes and our subset of 665 genotypes with available resequencing. Another hypothesis is that genotypic variation at *SbWRKY86* is not causal for *RMES2* HPR but *trans*-regulating elements modulate this transcription factor (Atamian *et al*., 2012; Li *et al*., 2015). Finally, it is possible that *SbWRKY86* is not involved in *RMES2* resistance and one or both of the QTL (S09_61521444, S09_62669680) are in higher LD with the causal gene(s). This QTL is in a gene dense region of Chr09 with 278 genes annotated between 61–63 Mb, demonstrating the need for fine-mapping or variant testing to confirm or exclude the hypothesis that *SbWRKY86* underlies *RMES2* resistance.

The peak *RMES2* variant was at high frequency in global germplasm as well as dispersed across Africa (Figure 3e). The resistant allele’s high frequency in historic breeding germplasm and sorghum conversion lines indicates *RMES2* has been present in North America for over half a century and is widely available for breeding in diverse and adapted backgrounds (Stephens *et al*., 1967). This is promising for breeders seeking additional HPR for cultivar development and will accelerate deployment when markers become available. *RMES2* KASP [kompetitive allele-specific polymerase chain reaction (PCR)] marker development is currently underway.

### *RMES2* bolsters *RMES1* to improve global HPR durability

The Chibas breeding program at the University of Quisqueya in Haiti breeds mixed-use temperate sorghums in high aphid pressure environments (Muleta *et al*., 2022). A breeding scheme which maximizes recombination with globally diverse founders of the breeding population provided strong statistical power to identify novel resistance associations that may be undetectable in association panels (e.g. *RMES1*) (Cockram and Mackay, 2018). *RMES1* and *RMES2* were the most significant QTL for mid-season resistance, however *RMES1* was not associated with resistance at early resistance (Figure 4b,c). One hypothesis is that there were abiotic (nutrient, environmental) or biotic factors other than *M. sorghi* pressure determining fitness at early stages different from late developmental stages. Another hypothesis is that the *RMES1* phenotype is developmentally-regulated akin to adult plant resistance, however, antibiosis-resistance was seen in the seedling stage in controlled greenhouse experiments (Figure 2a,b) and *RMES1* was first mapped in a biparental population at seedling stage in controlled settings (Wang *et al*., 2013). It is also possible that *RMES1* resistance is present throughout development where its relative HPR contribution increases with maturity but is undetectable in early-season field environments.

A common HPR source for early and late resistance would be expected to have positive selection and would indicate collective benefit to resistance. Agreeing with this hypothesis, variants at *RMES2* show selection signatures (Figure 4a). The selection of *RMES2* with *RMES1*, their additive contribution to resistance in the Haitian breeding program, and the presence of *RMES2* in global and historic sorghum genotypes led us to conclude that *RMES2* is bolstering *RMES1* HPR durability globally in breeding programs where sorghum aphid resistance is a target trait.

### Dhurrin and other candidate sources of quantitative and seedling resistance

We found evidence of additional sources of resistance in the association panels when controlling for the major *RMES2* association and in globally-admixed F_2_ lines, particularly at seedling stage, that may be sources of quantitative or early-season HPR. The cyanogenic glucoside dhurrin is known to provide resistance to chewing insects and may be an aphid resistance mechanism, although not proven (Dreyer *et al*., 1981; Gruss *et al*., 2022). As a major target trait in forage breeding, variation for dhurrin biosynthesis and bioactivation have been well studied with major variation mapped to Chr01 and Chr08, respectively (Hayes 2015). The biosynthetic gene cluster (*CYPE71E1*, *CYPE79A1*, *UGT85B1*) coincided with associations on Chr01 for resistance confounded by RMES2 (Figure 3b) and early-season resistance (Figure 4b). Dhurrin content is higher in seedlings and biosynthesis genes are highly expressed through vegetative development, which may explain the lack of association for resistance at maturity (Figure 4c) (Gleadow 2021, Halkier and Moller 1989). An association with early-season resistance association on Chr08 is selected for, however it falls over 1 Mb away from the bioactivation gene cluster and may represent a non-cyanogenic mechanism (Figure 4b,a, Table 1,2). Further investigations are warranted of dhurrin metabolism and catabolism as a mechanism of resistance against sorghum aphid.

Epicuticular waxes are known to affect aphid infestation as well as other biotic and abiotic traits in sorghum (Premachandra 1994, Weibel 1986). Mutant genotypes lacking waxy depositions, such as *bloomless2* which alters monoacyl glycerol and 32-C-alcohol wax profiles, are preferred for sorghum aphid feeding compared to wild type sorghum but does not affect aphid reproduction (Cardona 2022). Despite the established relationship between bloomless mutants and aphid HPR, natural variation for waxes contributing HPR has not been reported. Natural variation for epicuticular wax in an association panel was identified on Chr03 (∼66.6 Mb) and within 0.5 Mb of the S03_66114610 association for early-season resistance (Figure 4b) (Elango 2020). Wax deposition in sorghum is believed to be pleiotropic, benefitting water use efficiency traits, reducing forage digestibility, and preventing foliar pathogens (Cummins and Sudweeks, 1976; Jenks *et al*., 1994; Premachandra *et al*., 1994; Burow *et al*., 2008). Additional pleiotropy for sorghum aphid resistance would need to be considered in using this as a source of HPR.

### Next steps to mechanisms and global durability

Here we elucidated antibiosis-resistance provided by *RMES1* which places selection pressure on sorghum aphid populations, however, additional sources of resistance including *RMES2* are increasing global durability of HPR. Genomic and functional characterization of both *RMES1* and *RMES2*, as well as validation of dhurrin as an HPR source, will improve the molecular understanding of the genotype to phenotype map and inform breeding strategies. Global agricultural resilience will benefit from sustainable sorghum production that is durably aphid-resistant as climate change and anthropogenic factors continue to challenge food security around the world.

## ABBREVIATIONS

HPR: Host plant resistance
*RMES1*: *Resistance to Melanaphis sorghi 1*
NIL: Near-isogenic line
SNP: Single nucleotide polymorphism
GLM: General linear model

## ACKNOWLEDGEMENTS

This study is made possible by the support of the American People provided to the Feed the Future Innovation Lab for Collaborative Research on Sorghum and Millet through the United States Agency for International Development (USAID) under Cooperative Agreement no. AID-OAA-A-13-00047. The contents are the sole responsibility of the authors and do not necessarily reflect the views of USAID or the United States Government. Phenotyping and evaluation also received additional support from the Haitian Ministry of Agriculture, Natural Resources, and Rural Development through (PITAG project supported by the Interamerican development bank).

## AUTHOR CONTRIBUTIONS

GM, GP, VN, BR, and CV: conceived research hypotheses. GM, GP, VN, and CV: designed experiments. TF: NIL development. CV: Genomic and phenotypic characterization of NILs. BR and CV: genome-wide association studies. JRC and GP: breeding population development and phenotyping. CV and GM: manuscript development.

## SUPPLEMENTAL MATERIALS

**Supplemental Data 1**

Top variants in combined association panels and their associations with resistance phenotype determined by GLM

**Supplemental Data 2**

Top variants in combined association panels and their associations with resistance phenotype determined by GLM with *RMES2* included as a fixed covariate

**Supplemental Data 3**

Global sorghum accessions with metadata (longitude, latitude, germplasm origin) used for global and breeding frequency analyses

**Supplemental Data 4**

Top variants in globally admixed breeding population segregating for resistance and their associations with early-season survival

**Supplemental Data 5**

Top variants in globally admixed breeding population segregating for resistance and their associations with mid-season survival

## DATA AVAILABILITY

Raw whole-genome sequencing reads, association panel genotype data, Haitian genotype data, and geographic distribution data have been deposited at Dryad Digital Repository (https://doi.org/10.5061/dryad.rv15dv4f6).

